# Visualization of two architectures in class-II CAP-dependent transcription activation

**DOI:** 10.1101/2019.12.13.875658

**Authors:** Wei Shi, Yanan Jiang, Yibin Deng, Zigang Dong, Bin Liu

## Abstract

Transcription activation by cyclic AMP receptor protein (CAP) is the classic paradigm of transcription regulation in bacteria. CAP was suggested to activate transcription on class-II promoters via a recruitment and isomerization mechanism. However, whether and how it modifies RNA polymerase (RNAP) to initiate transcription remains unclear. Here we report cryo-EM structures of an intact *E. coli* class-II CAP-dependent transcription activation complex (TAC) with and without *de novo* RNA transcript. The structures reveal two distinct architectures of TAC and show that CAP-binding induces substantial conformational changes in all the subunits of RNAP and consequently widens the main cleft of RNAP considerably to facilitate DNA promoter entering and formation of initiation open complex. These structural changes vanish during further RNA transcript synthesis. The observations in this study suggest a unique activation mechanism on class-II promoters that CAP activates transcription by first remodeling RNAP conformation and then stabilizing initiation complex.

## Introduction

Transcription, the first step of gene expression, is regulated by various transcription factors. How these transcription factors modulate RNA polymerase (RNAP) is significant for understanding the mechanisms of transcription and gene regulation. Cyclic AMP (cAMP) receptor protein (CAP) is a classic dimeric global transcription factor in bacteria, which is activated by its allosteric effector cAMP [1–3]. CAP is able to activate and initiate transcription on more than 100 promoters in *E. coli* [1], mainly via two different mechanisms: class-I and class-II, according to its interaction mode with RNAP holoenzyme [4–6], the main enzyme comprising of a five-subunit core enzyme (α_2_ββ′ω) and a sigma factor and being responsible for RNA synthesis in cells [7,8]. On class-I promoters, such as *lac* promoter, CAP binds at the −61.5 site of the promoter and interacts with RNAP holoenzyme by the carboxyl-terminal domain of the alpha subunit (αCTD) of RNAP and activates transcription via a recruitment mechanism [9,10]. On class-II promoters, as exampled by *gal* promoter, CAP binds at the −41.5 site of the promoter and makes contact with multiple subunits of RNAP holoenzyme and initiates transcription via a recruitment and isomerization mechanism [4,11–14].

Our recent cryo-electron microscopy (cryo-EM) structure of the *E. coli* class-I CAP-dependent transcription activation complex (CAP-TAC) [15] has shown the overall architecture of the class-I complex and proven the recruitment mechanism via simple stabilization of initiation complex on class-I promoters. Recent crystal structure of the *Thermus Thermophilus* (*T.th*) class-II TAP (a CAP homolog in *T.th*)-dependent TAC (TAP-TAC) [16] has also displayed an architecture in which there are no apparent CAP binding-induced conformational changes in RNAP, and consequently suggested a simple stabilization mechanism for class-II activation as well. However, considering the extensive contact interface between CAP dimer and RNAP holoenzyme in the class-II TAC, it is possible that there are the CAP binding-induced conformational changes existing in RNAP, which are difficult to be captured by X-ray crystallography due to crystal packing, but accessible in electron microscopy. Thus, the need of investigating the mechanism of class-II activation still remains.

In this study, we have determined three cryo-EM structures of the intact *E. coli* class-II CAP-TAC at around 4.2-4.5 Å resolutions, containing a CAP dimer, a σ^70^-RNAP holoenzyme, a complete class-II CAP-dependent promoter DNA, and with or without a *de novo* synthesized RNA oligonucleotide, and also revealed two distinct architectures of the class-II TACs: state 1 and state 2. The state 1 architecture visualizes the intermediate activation complex, suggesting that the CAP binding-induced major conformational changes in RNAP lead to a result of remarkable widening of the main cleft, which would greatly facilitate DNA prompter entering into it and further isomerizing to transcription open complex. On the basis of these structures, we suggested the activation mechanism on class-II promoters that CAP activates transcription by first remodeling RNAP conformation and then stabilizing initiation complex.

## Results and Discussion

### Overall structures of the *E. coli* class-II CAP-TACs without RNA transcript

To investigate whether and how CAP-binding induces conformational changes of RNAP to activate transcription on class-II promoters, a synthetic DNA scaffold representing the complete class-II CAP-dependent promoter (from −64 to +14, 78-bp in total) as observed in the *gal* promoter (Fig 1A) was designed to assemble the intact *E. coli* class-II CAP-dependent TAC (CAP-TAC) *in vitro* (Fig EV1). To test the effect of RNA transcript synthesis on the overall architectures, the complex was also assembled in the presence of the nucleotides (ATP and GTP) to synthesize RNA transcript.

**Figure 1.**
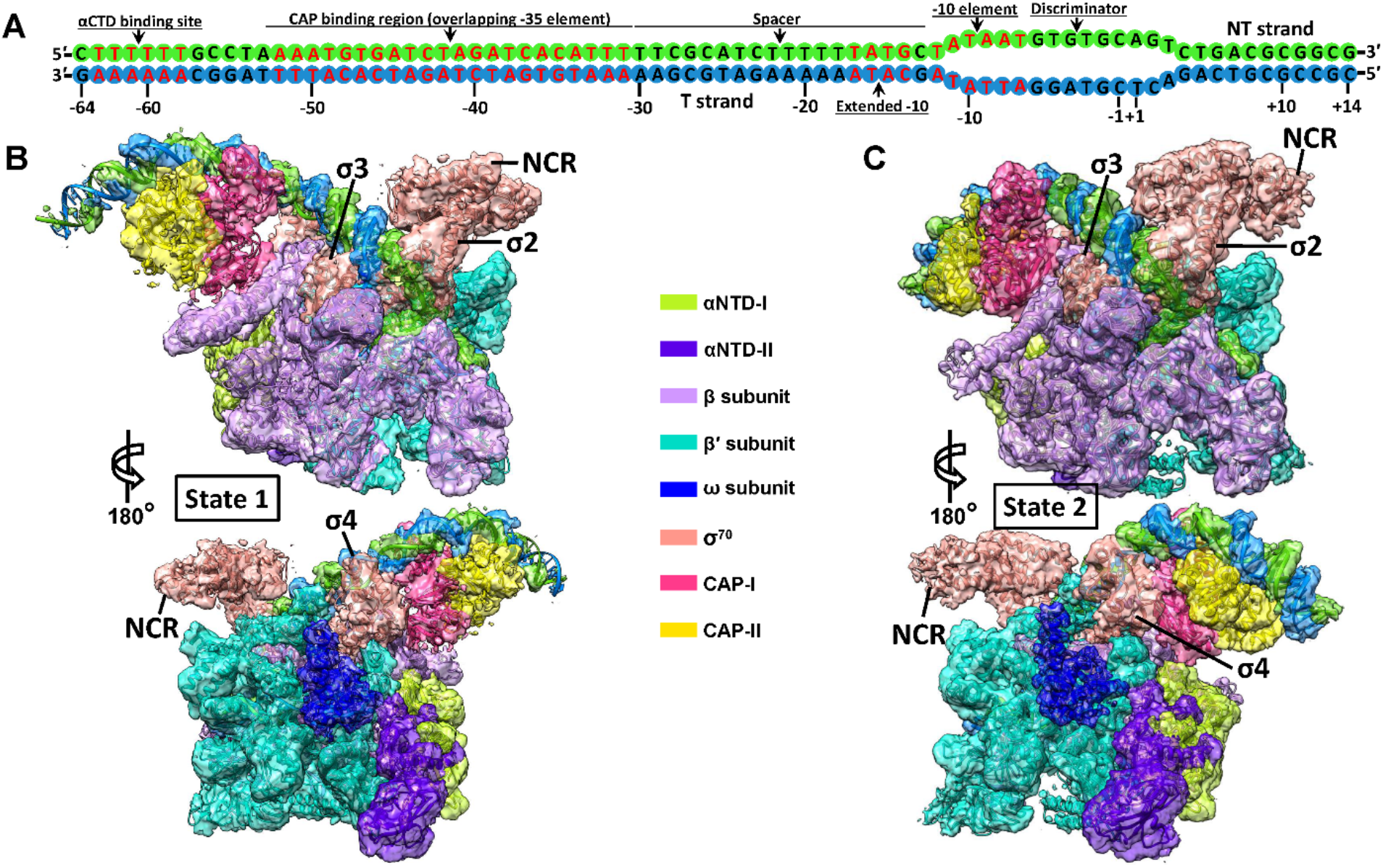
Cryo-EM reconstructions of the class-II CAP-TAC without RNA transcript. A Schematic representation of the synthetic promoter DNA scaffold (78-bp) in the class-II CAP-TAC. B, C Overviews of the cryo-EM reconstruction maps of the *E. coli* class-II CAP-TAC without RNA transcript at 4.52 Å (B, state 1) and 4.29 Å (C, state 2) resolution, respectively. The individually colored density maps, created by color zone and split in Chimera and shown in a contour of 8 root-mean-square (RMS), are displayed in transparent surface representation to allow visualization of all the components of the complex.

The isolated complexes were subjected to cryo-EM analyses. The cryo-EM single-particle reconstructions of the intact class-II CAP-TAC without NTP incubation showed two different architectures of the structure: state 1 and state 2 at overall resolutions of 4.29 Å and 4.19 Å, respectively (Fig EV2; Table EV1). In the cryo-EM maps, the densities for RNAP holoenzyme allowed the unambiguous docking of its components, while the CAP binding region is poorly-defined due to high structural flexibility. The αCTD of RNAP is invisible in density maps although a specific DNA sequence for its binding is included in the DNA scaffold. The non-conserved region (NCR) of σ^70^ and the ω subunit of the holoenzyme are visible at a low contour level, owing to their structural flexibilities and low occupancies. Further focused classifications and refinements with a sphere covering only the CAP binding region improved the density for this portion and generated final overall 4.52 Å and 4.29 Å resolution maps for the state 1 and state 2 structures, respectively (Fig 1B and C).

### Two distinct architectures reveal CAP-binding induced conformational changes in RNAP

The structures of state 1 and state 2 CAP-TACs without NTP incubation show the formation of a transcription open complex (RPo). Remarkably, the state 1 structure of CAP-TAC exhibits a distinct architecture in RNAP from the one observed in the previous cryo-EM structure of *E. coli* RPo in the absence of CAP binding [17] (Fig 2A). Compared to the previous RPo structure, all the subunits of RNAP in the state 1 structure display substantial conformational changes, especially the domains around the main cleft (Fig 2B). The secondary structures in the β-flap, protrusion, lobe, sequence insertion 1 (SI1), and SI2 domains of the β subunit of RNAP shift to a maximum distance of 7.5 Å, 14.4 Å, 8 Å, 12 Å, and 26 Å, respectively. The movements of those in the clamp, jaw, and SI3 domains of the β′ subunit of RNAP reach to a maximum of 12 Å, 7 Å, and 9 Å in distance, respectively. These conformational changes lead to a result of remarkable widening of the main cleft accommodating DNA promoter, which would greatly facilitate DNA prompter entering into the cleft and further isomerizing to transcription open complex (Fig 2B). Since the major difference between the two complexes is whether the CAP dimer is present or not, these significant conformational changes observed in the structure of state 1 CAP-TAC should result from CAP binding.

**Figure 2.**
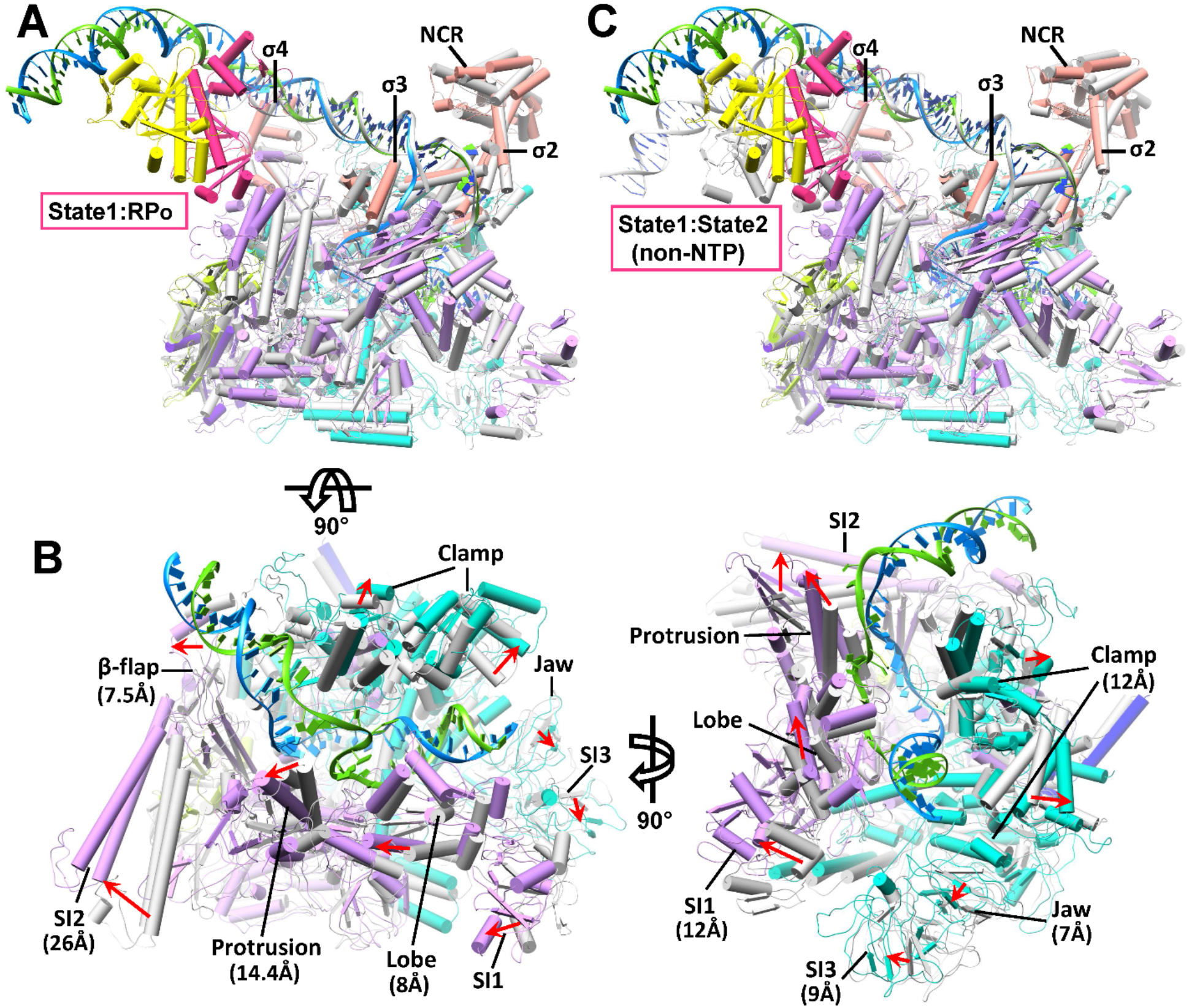
The CAP binding-induced conformational changes in RNAP. A Superimposition of the state 1 CAP-TAC (colored) with the previous transcription open complex (RPo) (gray, PDB 6CA0) [17] via the σ2 and σ3 domains of σ^70^. The components are shown in pipes and planks representation. The color scheme for the state 1 structure is same as that in Figure 1. B The close-up views of the main cleft along two directions. The DNA promoter from RPo and all the σ^70^ proteins are omitted for clear representation. The movement directions and maximal distances of the secondary structures in the domains surrounding the main cleft are labeled using red arrows and specific values, respectively. C Superimposition between the state 1 (colored) and state 2 CAP-TAC (gray) without RNA transcript via the σ2 and σ3 domains of σ^70^.

Previous biochemical study has shown that in the class-II activation, the interaction between the activating region 2 (AR2) of CAP and the amino-terminal domain of the alpha subunit (αNTD) of RNAP would increase the rate constant for isomerization from the closed to open complexes and therefore suggested AR2-αNTD interaction would either trigger an allosteric change of RNAP or stabilize the intermediate states between the closed and open complexes [11]. Our observation actually provided structural support for the first possibility. Clamp dynamics is significant in facilitating DNA template melting [18,19]. The opening and close of the clamp domain were apparently observed in prokaryotic and eukaryotic RNAP complexes [20–23]. The two TAC structures (state 1 and state 2) were superimposed with the previous *Mycobacterium tuberculosis* (*Mtb*) RNAP-DNA complexes (PDB 6EDT, 6EEC, 6EE8, and 6BZO), and the results showed that the main cleft in our state 2 architecture displays a similar narrow status as the one in the clamp-closed RPo structure (PDB 6EDT), while the cleft in the state 1 structure is as wide as that in the clamp-open *Mtb* RNAP-DNA complex (PDB 6BZO) (Fig EV3A-C). Similarly, a contraction and expansion of the main cleft in eukaryotic RNAP I has been reported as well [24]. In summary, all these evidences have supported the presence of the class-II TAC with state 1 architecture observed in this study.

In addition to the unique architecture in the RNAP portion, the relative position of CAP dimer and its bound DNA in the structure of state 1 CAP-TAC, suggested by the density in the cryo-EM map (Figs 1B and EV4A), is different from that shown in the state 2 CAP-TAC structure. There is an around 60° switch between the two positions of the CAP-bound DNA (Fig EV4B). This new position of the CAP dimer in state 1 apparently extends the contact interface between it and the β subunit of RNAP. Consequently, the SI2 of the β subunit rotates and shifts a huge distance (26 Å in maximum) to interact with CAP (Fig 2B). The β protrusion, β lobe, β′ jaw, and β′ SI3 domains, which all surround the main cleft, therefore make movement due to pulling force transduced through interactions among these domains. The driving force modulating the β′ clamp domain could be transferred through the σ4 domain or DNA promoter. The RNAP portion of the state 2 CAP-TAC shares a similar architecture as that observed in the previous RPo structure [17], and therefore comparison between the structures of the state 1 and state 2 CAP-TACs also demonstrates the mentioned considerable structural inconsistence in RNAP (Fig 2C).

### The conserved interactions in the TACs

The structure of state 2 CAP-TAC without RNA transcript defined the interactions among CAP, promoter DNA, and RNAP holoenzyme as well (Fig EV5A and B). The positively charged residue (K100) in the AR2 of CAP is close to the negatively charged residues (D164 and E165) in the 165 determinant of the αNTD of RNAP (Fig EV5C). The interface between the AR2 of CAP and the β-flap domain of RNAP is the main contact region, primarily involving hydrogen bonds (Fig EV5C). The AR3 of CAP makes interactions with the 596 determinant in the σ4 domain of σ^70^ mainly by salt bridges and hydrogen bonds, as well as with the β-flap tip (β-FT) by hydrogen bonds. These observed interactions stabilize the activation complex and are consistent with those shown in the previous structural and mutagenesis studies [11,12,16,25,26].

### *De novo* RNA transcript synthesis promotes the conversion of the state 1 to state 2

The cryo-EM density map for the complete class-II CAP-TAC with a *de novo* RNA transcript was reconstructed at an overall 4.0 Å resolution (Fig EV6; Table EV1). Further focused classifications and refinements also improved the density for the CAP binding region and generated a final overall 4.35 Å resolution map allowing us to confidently dock this region (Fig 3A). The αCTD is invisible in the density map as well, reconfirming the flexibility of this domain in the complex. The density at the active site suggests that a *de novo* synthesized RNA trinucleotide (GAG) starting from the - 1 position is present (Fig 3B), which is consistent with our previous structural studies [15,27].

**Figure 3.**
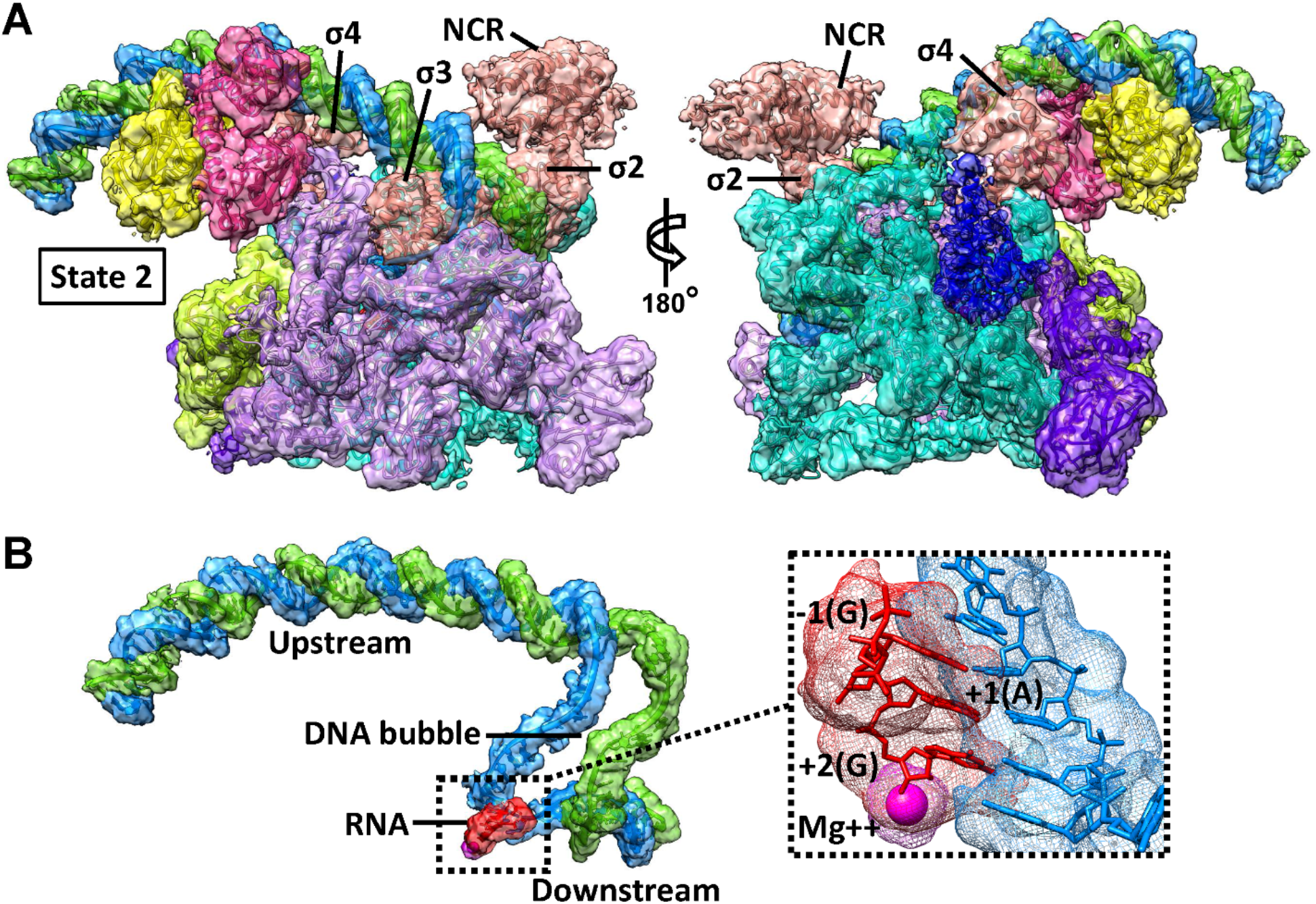
Cryo-EM reconstruction of the class-II CAP-TAC with *de novo* RNA transcript. A Overview of the cryo-EM reconstruction map of the *E. coli* class-II CAP-TAC with RNA transcript at 4.35 Å resolution and the state 2 architecture. The color schemes for the split density maps (8 RMS) and the docked components are same as in Figure 1. B A close-up view of the promoter DNA scaffold in the complex. The insert is the zoom-in view of the DNA-RNA hybrid region with the magenta Mg^2+^ sphere. A *de novo* synthesized RNA transcript (3-nt) starting from the −1 position with a GTP residue is displayed.

The structure of class-II CAP-TAC with *de novo* synthesized RNA transcript does not capture the state 1 architecture, but displays an architecture representing the state 2, and is highly similar to the one of CAP-TAC without NTP incubation (Fig EV7). These observations suggest that *de novo* RNA synthesis does not affect the overall architecture of the state 2 complex, but promotes the conversion of the state 1 to state 2. Subsequent to the clamp opening which allows DNA to be loaded into in the RNAP active center cleft, promoter unwinding and initial RNA synthesis trigger clamp closure, accounting for the high stability of initiation complexes and the high stability and processivity of elongation complexes [22]. Therefore, it is plausible that the architecture of the state 2 is more stable and resistible than that of the state 1 while resisting DNA scrunch generated during transcription initiation [28,29], and for that reason, no structure representing the state 1 was observed during RNA transcript synthesis. The overall architecture of the state 2 CAP-TAC is also similar to the structure of the *T.th* class-II TAP-TAC [16] (Fig EV8). No significant difference in the width of the main cleft is observed when superimposing them each other. However, the relative orientations of the CAP dimer and the domains on the interfaces between CAP and RNAP holoenzyme apparently shift, probably due to a longer DNA promoter containing the αCTD binding site used in this study.

## Concluding remarks

Our structural analyses of intact class-II CAP-TAC in the presence or absence of *de novo* RNA transcript synthesis displayed three structures with two distinct architectures. In the state 1 architecture, the CAP dimer with the bound promoter DNA swings ~60° from the putative position to make more extensive contacts with the β subunit of RNAP including SI2, and therefore induces major conformational changes in all the subunits of RNAP and significantly widens the main cleft; while in the state 2 architecture, the CAP dimer binds and bends the promoter DNA, and also interacts with the αNTD, β-flap, and σ4 domains as shown previously, but does not remodel the main cleft. These observations provide insights into how CAP binding induces the conformational changes of RNAP and activates transcription on the class-II DNA promoters (Fig 4). Based on our structures, it is presumable that once CAP dimer binds to RNAP holoenzyme and promoter DNA, they first construct a complex with the state 1 architecture in which the CAP binding-induced conformational changes in RNAP greatly widen the main cleft to facilitate DNA promoter enter into it and further form an initiation open complex. Partial of the complexes with the state 1 architecture would transfer to the ones at the state 2 with the formation of initiation open complex. With the incorporation of NTP, all the complexes at the state 1 convert to those with the state 2 architecture since the state 2 is more resistible for the initiation stage with DNA scrunch happening. Thus, on the class-II promoters, CAP activates transcription by first remodeling RNAP conformation and then simply stabilizing initiation complex. This novel transcription activation mechanism in bacteria, although needs further investigations to examine, would be, as of yet, unique for understanding gene regulation in prokaryotic system.

**Figure 4.**
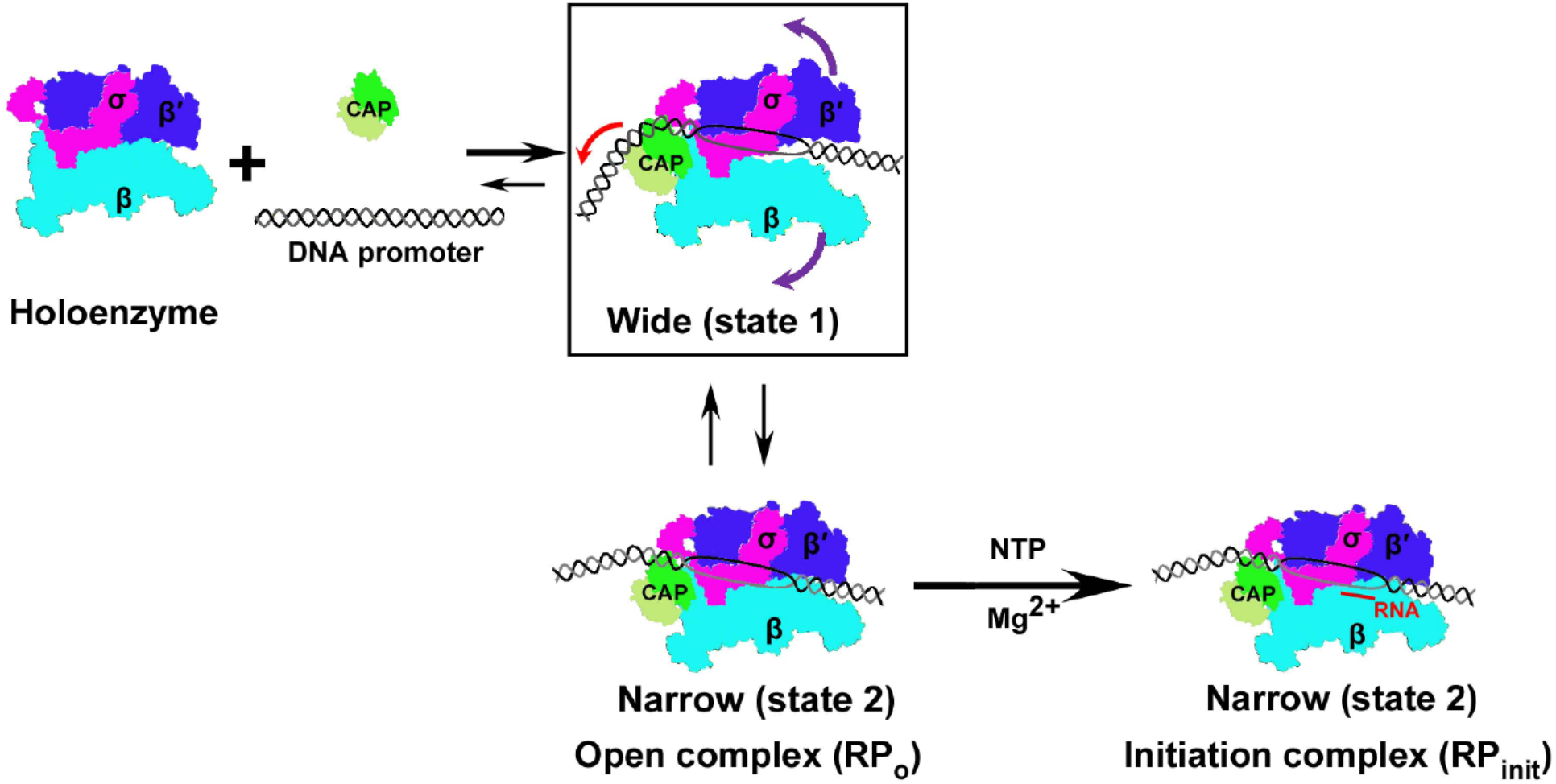
The mechanism of transcription activation on class-II promoters. A schematic cartoon model of CAP activating transcription on class-II DNA promoters is presented. When the CAP dimer interacts with RNAP holoenzyme and the class-II DNA promoter, it would first induce conformational changes in RNAP and consequently significantly widen the main cleft to form a CAP-TAC with the state 1 architecture, which facilitates the DNA promoter entering into the main cleft and further forming a transcription open complex. The complex at the state 1 can convert to the one with the state 2 architecture that contains narrow main cleft with the formation of initiation open complex. With transcription initiation and the synthesis of RNA transcript, all the complexes with different states would convert to the ones with the state 2 architecture. The colored arrows in the rectangle indicate the individual movement directions.

## Materials and Methods

### Preparation and assembly of *E. coli* class-II CAP-TAC

*E. coli* RNAP, σ^70^, and CAP proteins were expressed and purified as described [15,27,30,31]. RNAP was then mixed with an excess of σ^70^ and loaded onto a 16/60 Superdex 200 prep grade gel filtration column (GE Healthcare) with buffer A (20 mM TRIS pH 8.0, 50 mM NaCl). Fractions containing the σ^70^-RNAP holoenzyme were pooled and concentrated to around 10 mg ml^-1^. To construct a class-II CAP-dependent transcription activation complex, we used a synthetic DNA scaffold corresponding to the promoter region between positions −64 and +14 relative to the expected transcription start site (Fig 1A). The synthetic promoter, which contains the αCTD binding site, the CAP protein binding region overlapping the −35 element, the extended −10 motif, and the consensus −10 element, was prepared by annealing the non-template strand to an equal molar amount of the template strand DNA that is complementary to the NT-strand except for a 6-nucleotide (nt) discriminator region. The class-II CAP-TACs were assembled by directly incubating the σ^70^-RNAP holoenzyme with a two-fold molar excess of the pre-formed DNA promoter and CAP protein in buffer B (20 mM TRIS pH 7.5, 50 mM NaCl, 5 mM MgCl_2_) at 37°C for 10 minutes in the presence of 0.2 mM cyclic AMP (cAMP), and with or without ATP and GTP (0.2 mM each).

### Cryo-EM sample preparation and data acquisition

The incubated samples were re-purified by a Superdex 200 Increase 10/300 GL gel filtration column (GE Healthcare) with buffer B to remove the extra CAP protein and nucleic acids. A droplet of 3.5 μl of the re-purified class-II CAP-TACs at 1 µM was placed on Quantifoil 1.2/1.3 300 mesh Cu grids (EM Sciences) glow-discharged at 15 mA for 60 sec. The grid was then blotted for 3 sec at 4°C under conditions of 100% humidity, and flash-frozen in liquid ethane using a Vitrobot mark IV (FEI). Cryo-EM data were collected on a 300 kV Titan Krios microscope (FEI) at The Hormel Institute equipped with a Falcon III direct electron detector (FEI). A 100 μm objective aperture was applied during data collection. Micrographs were recorded in the counted mode at a pixel size of 0.9 Å, with the defocus ranging from −0.8 to −2.6 μm. Recorded at a dose rate of 0.8 e-/pixel/sec (1 e^−^/Å^2^/sec), each micrograph consisted of 30 dose-framed fractions. Each fraction was exposed for 1.5 (CAP-TAC without NTP incubation) or 1.2 (CAP-TAC with NTP incubation) sec, resulting in a total exposure time of 45 sec (CAP-TAC without NTP incubation) or 36 sec (CAP-TAC with NTP incubation) and total dose of 45 or 36 e^−^/Å^2^. Fully automated data collection was carried out using EPU (FEI).

### Image processing

Data was processed using cisTEM [32]. A total of 2,369 (CAP-TAC without NTP incubation) or 2,819 (CAP-TAC with NTP incubation) movies were collected. Beam induced motion and physical drift were corrected followed by dose-weighing using the Unblur algorithm [33]. The contrast transfer functions (CTFs) of the summed micrographs were determined using CTFFIND4 [34]. Particles were then automatically selected based on a matched filter blob approach with the parameters: maximum particle radius (80 Å), characteristic particle radius (60 Å), threshold peak height (1.5, standard deviation above noise), 30 Å of highest resolution used in picking, and avoiding high variance areas and areas with abnormal local mean [35]. Initially, 477,218 particles (CAP-TAC without NTP incubation) or 628,208 particles (CAP-TAC with NTP incubation) were picked and extracted to construct a refinement package with 200 Å of largest dimension and 384 pixels of stack box size. 2D classifications [36] were performed using 300-40/8 Å (start/finish, high-resolution limit) data and always without inputting starting references.

For the data of the sample CAP-TAC without NTP incubation, 103 of 200 classes in the first round of 2D classification were manually chosen by discarding the ones containing unsuitable particles and obvious Einstein-from-noise to construct a new refinement package (215,253 particles) and subjected to the second round of 2D classification.

Then, 33 of 200 good classes were picked to construct a new refinement package with 84,537 particles and subjected to Ab-Initio 3D reconstruction [37] to generate an initial 3D model using 20-8 Å resolution data. This initial 3D model was set as the starting reference for the further 3D Auto refinement. Fourier Shell Correlation at a criteria of 0.143 reported a 4.29 Å resolution for the map outputted from Auto refinement [38]. The flexible and disordered CAP-binding region urged us to perform a focused classification (3 classes) and refinement using 300-10 Å resolution data and a sphere, which has a 60 Å radius and covers the CAP-binding region, in 3D manual and local refinement. Good classes were obtained after 17 cycles of refinements. The second class (46.42%) was reconstructed on 39,242 particles, suggesting an architecture of state 1 with 4.4 Å resolution, in which the architecture of RNAP core and the locations of CAP dimer and its bound DNA are distinct from the one that was previously reported [16,17]. However, there still has minor density in the previously displayed location of CAP dimer. Class 1 shows a similar state at 6.12 Å resolution, while much more mixed states are present in the CAP-binding region of the map from class 3. Further focused classification and refinement with the classes from the first round of focused classification generated clearer density map on the CAP binding region in class 2 and class We then combined the particles in this two classes to run Auto refinement and finally got a clearer density map with 4.52 Å resolution.

We also extended the selection range from the second round of 2D classification. 65 of 200 classes, including 115,593 particles, were chosen to subject to the third round of 2D classification. 36 of 80 classes (84,227 particles) were selected to generate a new 3D model (reverse handedness) by Ab-Initio 3D reconstruction. The new model was then refined to 4.19 Å using 3D Auto refinement, revealing an architecture of state 2. Further focused classification (3 classes) and refinement (10 cycles) with a 40 Å radius of sphere centered at the CAP-binding region and use of 300-10 Å resolution data generated much better density on the CAP-binding region from class 2 (37,456 particles, 44.47%, 4.29 Å).

For the data of the sample CAP-TAC with NTP incubation, 35 of 200 classes (171,254 particles) in the first round of 2D classification were manually chosen to subject to the second round of 2D classification. A final package with 109,437 particles (18 of 50 classes) was constructed to generate an initial 3D model using 20-8 Å resolution data by Ab-Initio 3D reconstruction. The model was refined to 4.14 Å using 3D Auto refinement and then 4 Å with further 3D manual and local refinement, revealing an architecture of state 2. Further focused classification (3 classes) and refinement (11 cycles) with a 60 Å radius of sphere centered at the CAP-binding region and use of 300-10 Å resolution data generated much clear density on the CAP-binding region from class 2 (33,455 particles, 30.57%, 4.35 Å).

### Structural modelling and refinement

The initial models were generated by docking the previous crystal or cryo-EM structures of the components into the individual cryo-EM density maps outputted from the focused classification and refinement using Chimera [39] and COOT [40], including *E. coli* RNAP core enzyme, the bubble region and the downstream DNA portion from the *E coli* σ^s^-TIC (PDB 5IPL) [27], and the upstream DNA portion and σ^70^ from the model of the class-I CAP-TAC (PDB 6B6H) [15]. The 4.35 Å cryo-EM density map from the sample of CAP-TAC with NTP incubation allowed us to build an RNA trinucleotide (GAG) at the active site starting from the −1 position. The omega subunits in all the three cryo-EM density maps are still docked according to the previous RNAP holoenzyme structures although poor densities suggest their flexibilities and low occupancies. The cryo-EM maps, representing the state 1 (4.52 Å) and state 2 (4.29 Å and 4.35 Å), from the focused classifications and refinements show clear density in the CAP-binding region, ensuring further docking of the whole CAP dimer domain using the models from the structures of the CAP-αCTD-DNA complex (PDB 1LB2) [41] and the class-I CAP-TAC (PDB 6B6H) [15]. In all the three cryo-EM maps, no density allowed us to dock αCTD although we have designed the DNA binding sequence for αCTD.

The intact models were then real-space refined using Phenix [42]. The final maps were put into a large P1 unit cell (*a* = *b* = *c* = 345.6 Å; α = β = γ = 90°) and structural factors were calculated in Phenix [42]. In the real-space refinement, minimization global and local grid search were performed with the secondary structure, rotamer, and Ramachandran restraints applied throughout the entire refinement. The final model has good stereochemistry by evaluation using MolProbity [43]. The local resolution maps were estimated and generated by ResMap [44] and MonoRes [45] using the online server (http://scipion.cnb.csic.es/m/myresmap). The statistics of cryo-EM data collection, 3D reconstruction and model refinement was shown in the Table EV1. All the figures were created using Chimera [39].

### Quantification and Statistical Analysis

Quantification, statistical analysis, and validation are implemented in the software packages used for 3D reconstruction and model refinement.

## Acknowledgements

We thank Y. Wang at Yale University for constructing *E. coli* CAP and σ^70^ plasmids. The cryo-EM data were collected at the cryo-electron microscopy facility in the Hormel Institute, University of Minnesota, which is funded by the Hormel Foundation. This work was supported by the starting-up funding granted to B.L. from the Hormel Institute, University of Minnesota.

## Author contributions

B.L. initiated and designed the experiments. W.S. performed protein sample preparations and assembly of the complexes used in the structure determination. W.S., Y.J. and B.L. performed cryo-EM grid preparation, screening, and optimization. B.L. conducted high throughput data collection on Titan Krios, image processing, map generation, and model building and refinement. W.S. Y.J. and B.L. performed structural analysis. W.S., Y.J., Y.D., Z.D., and B.L. wrote the manuscript.

## Conflict of interest

The authors declare no competing interests.

## Data Availability

The cryo-EM maps and the atomic coordinates generated in this study have been deposited in the Electron Microscopy Data Bank (EMDB) and the Protein Data Bank (PDB) under the accession numbers: EMD-20287 and 6PB5 (the class-II CAP-TAC without RNA transcript at the state 1), EMD-20288 and 6PB6 (the class-II CAP-TAC without RNA transcript at the state 2), and EMD-20286 and 6PB4 (the class-II CAP-TAC with RNA transcript at the state 2).

## Figure Legends

**Figure EV1.**
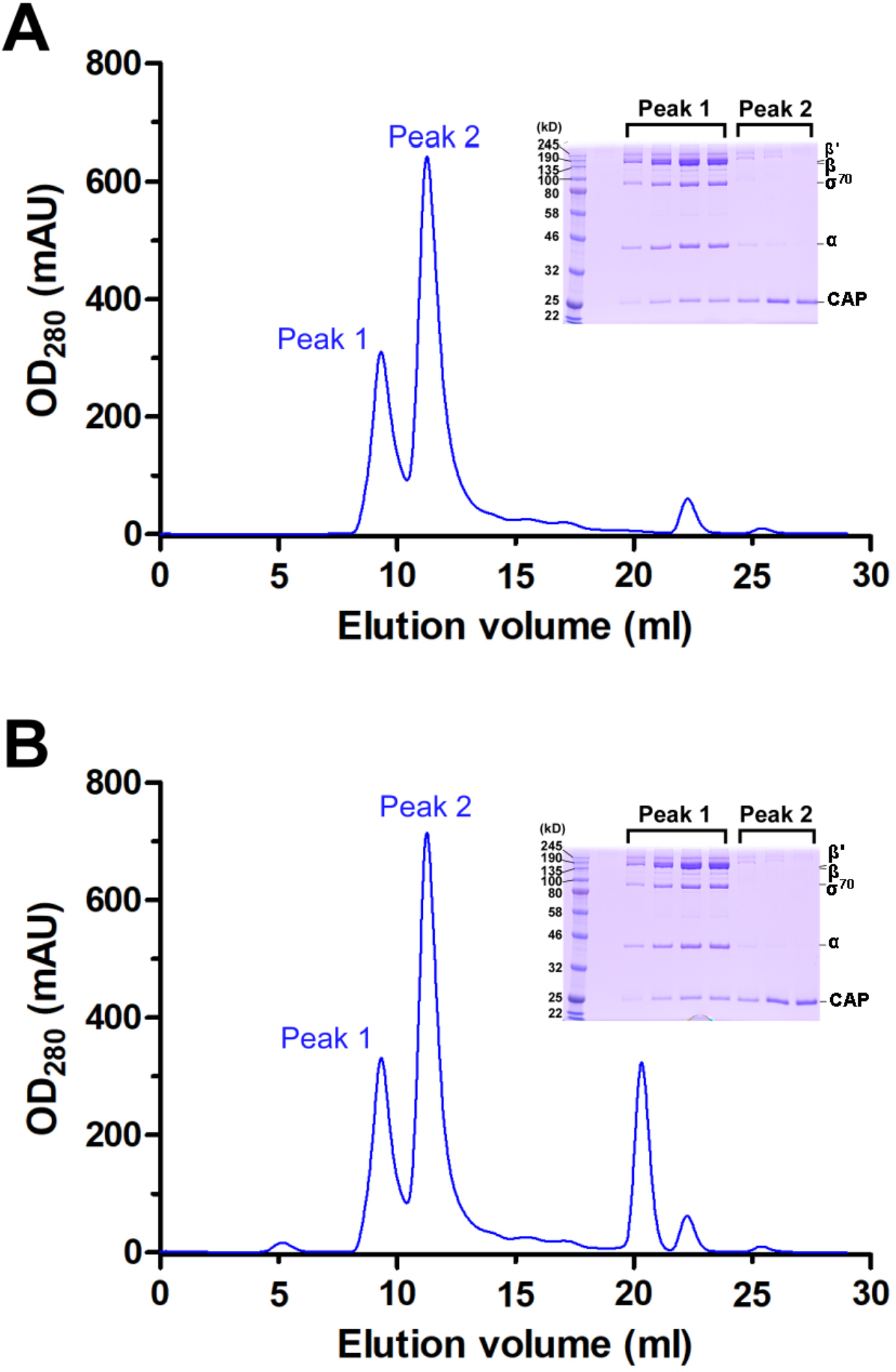
Isolation of the class-II CAP-TACs by size-exclusion chromatography. A The size-exclusion chromatography profile of the CAP-TAC without NTP incubation is presented. The inserted SDS-PAGE gel verified the presence of the complex. B The size-exclusion chromatography profile of the CAP-TAC with NTP incubation and the inserted SDS-PAGE gel visualizing the components are presented.

**Figure EV2.**
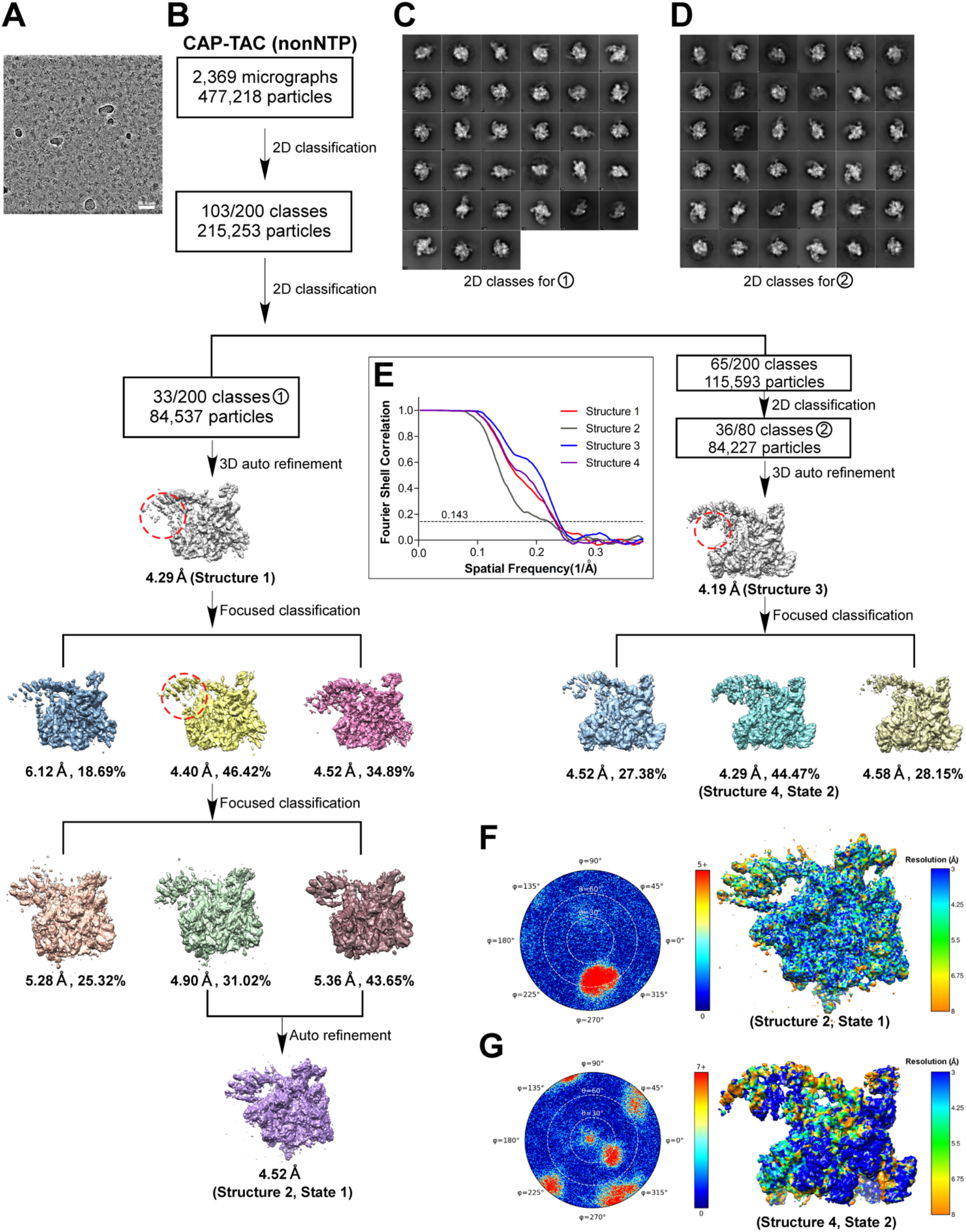
Cryo-EM images and data processing procedure for the CAP-TAC without RNA transcript. A A representative micrograph B Flow chart of the cryo-EM image processing (see Methods). C, D Selected 2D classes for structure 1 (C) and structure 3 (D), respectively. E Gold-standard Fourier Shell Correlations (FSCs) of the maps for structures 1-4. F Angular orientation distribution (left) of the particles used in the final reconstruction and local resolution map (right) for structure 2 (state 1). G Angular orientation distribution (left) of the particles used in the final reconstruction and local resolution map (right) for structure 4 (state 2).

**Figure EV3.**
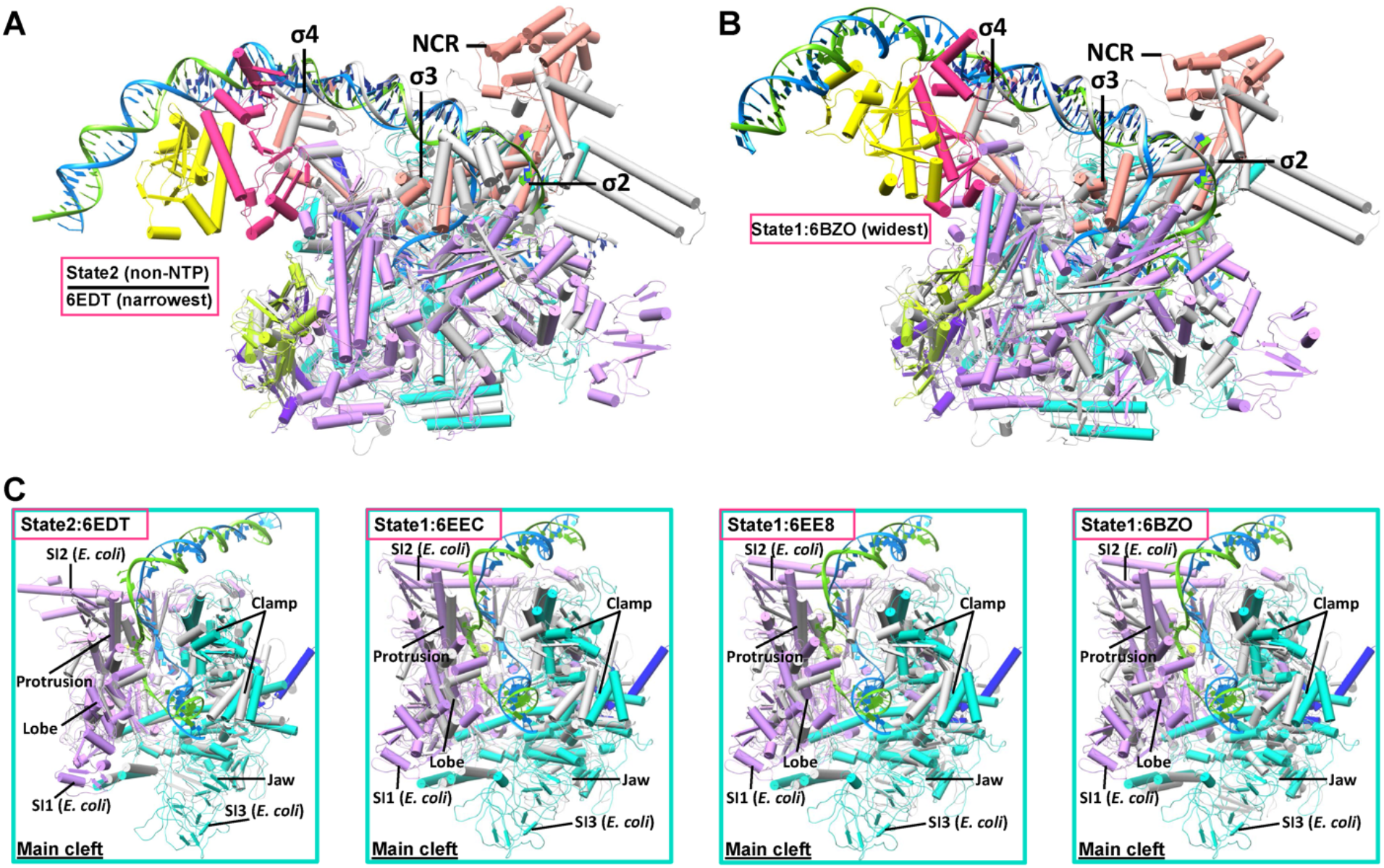
Superimpositions between *E. coli* class-II TACs and *Mtb* RNAP-DNA complexes. A, B Superimpositions of the *E. coli* class-II CAP-TAC (colored, state 2) with the *Mtb* RNAP-DNA-RbpA/CarD (gray, PDB 6EDT) and the *E. coli* class-II CAP-TAC (colored, state 1) with the *Mtb* RNAP-DNA-RbpA/Fidaxomicin (gray, PDB 6BZO) via the σ2 and σ3 domains are shown respectively. The color schemes are same as in Figure 1. 6EDT and 6BZO represent the one with the narrowest and widest cleft, respectively, in the determined *Mtb* RNAP-DNA complexes. C Zoom-in views of the superimpositions via the σ2 and σ3 domains (from left to right: state2:6EDT, state1:6EEC, state1:6EE8, state1:6BZO), in which 6EEC and 6EE8 represent the ones with the intermediate width of the main cleft. DNAs from TACs and RNAPs from all structures are shown. The results suggest that the observed opening and closing of the main cleft in this study are similar to those shown in the structures of the *Mtb* RNAP-DNA complexes.

**Figure EV4.**
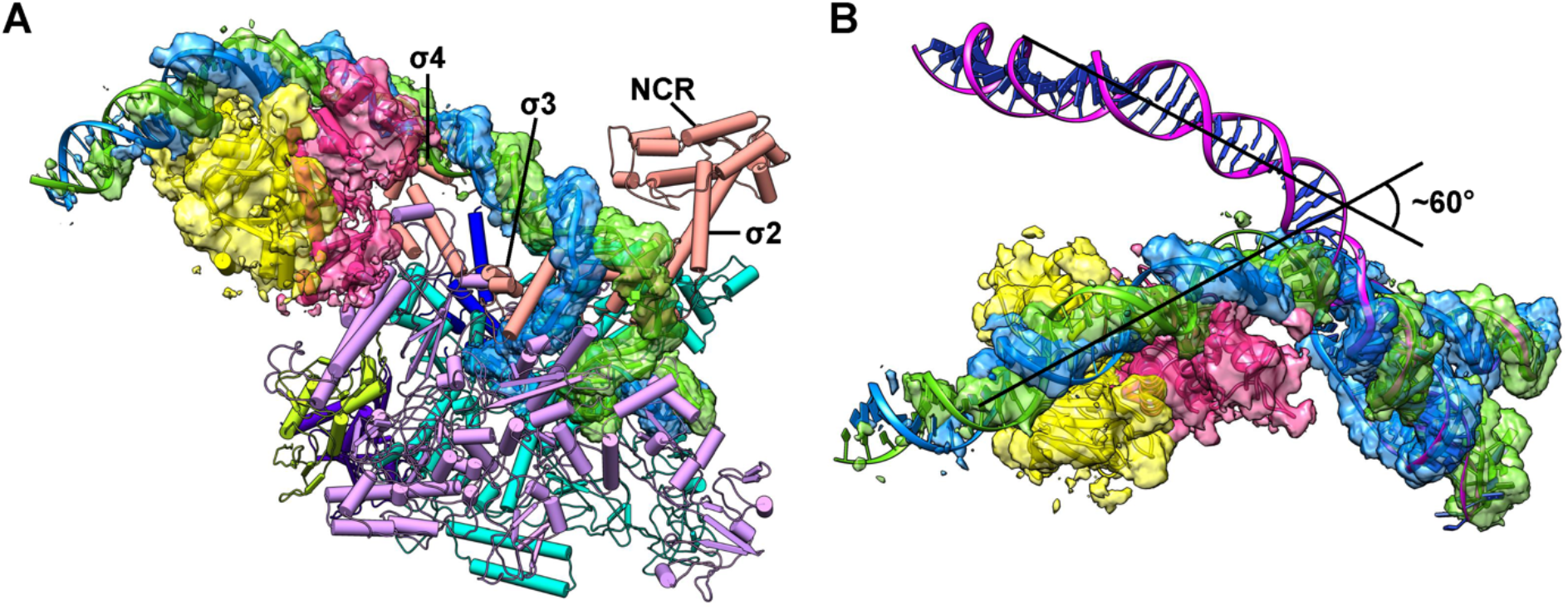
The new position of the CAP dimer and its bound promoter DNA in the CAP-TAC with the state 1 architecture. A The transparent split cryo-EM map for the CAP dimer region and the promoter DNA in the state 1 CAP-TAC is shown (contoured at 8 RMS). RNAP holoenzyme is displayed with pipes and planks representation. The color schemes are same as in Figure 1. B Superimposition of the promoter DNAs between the state 1 and state 2 (magenta) CAP-TACs without RNA transcript shows a ~60° swing between the two positions of the CAP dimer and its bound promoter DNA. The CAP dimer of state 2 and RNAP holoenzymes were omitted for clear representation.

**Figure EV5.**
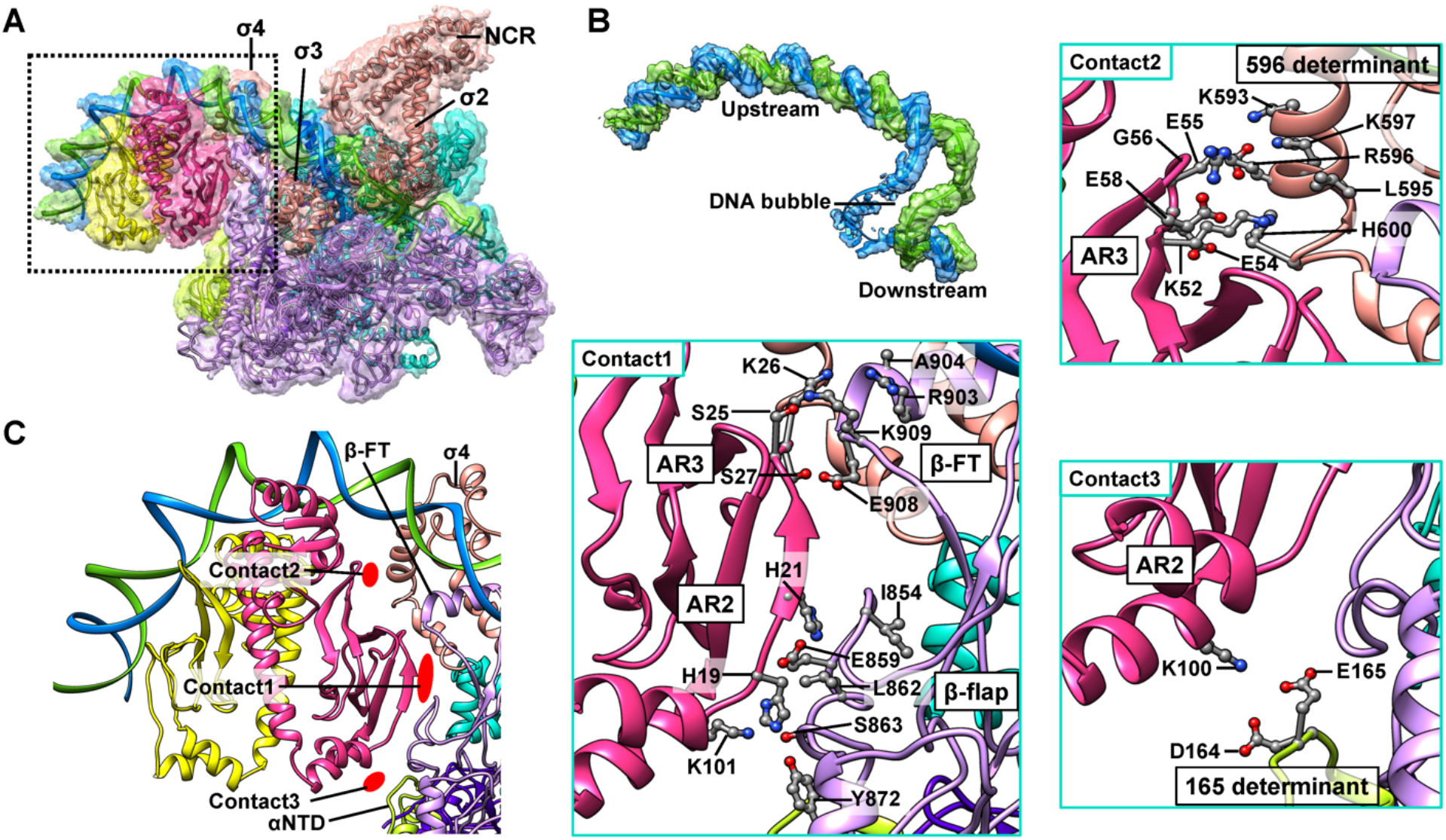
The interactions between CAP and RNAP holoenzyme in the state 2 CAP-TAC without RNA transcript. A Overview of the *E. coli* class-II CAP-TAC without RNA transcript. The transparent split cryo-EM maps (8 RMS) and the components are shown in the same color schemes as in Figure 1. B A close-up view of the promoter DNA in the complex. C A close-up view of the interface between CAP and RNAP holoenzyme and the zoom-in views of the three main contact areas. In the structure, the CAP dimer makes contacts with the 165 determinant (residues 163-165) of the αNTD and the β-flap domain (residues 854, 859, 862, 863, and 872) through its activating region R2 (AR2, residues 19, 21, 100, and 101), as well as with the β-flap tip (β-FT, residues 903-904, and 908-909) and the 596 determinant (residues 593-603) of σ4 using its AR3 (residues 25-27, and 52-58, respectively). It should be noted that there may be some shifts in the side chains of these residues at the current resolution.

**Figure EV6.**
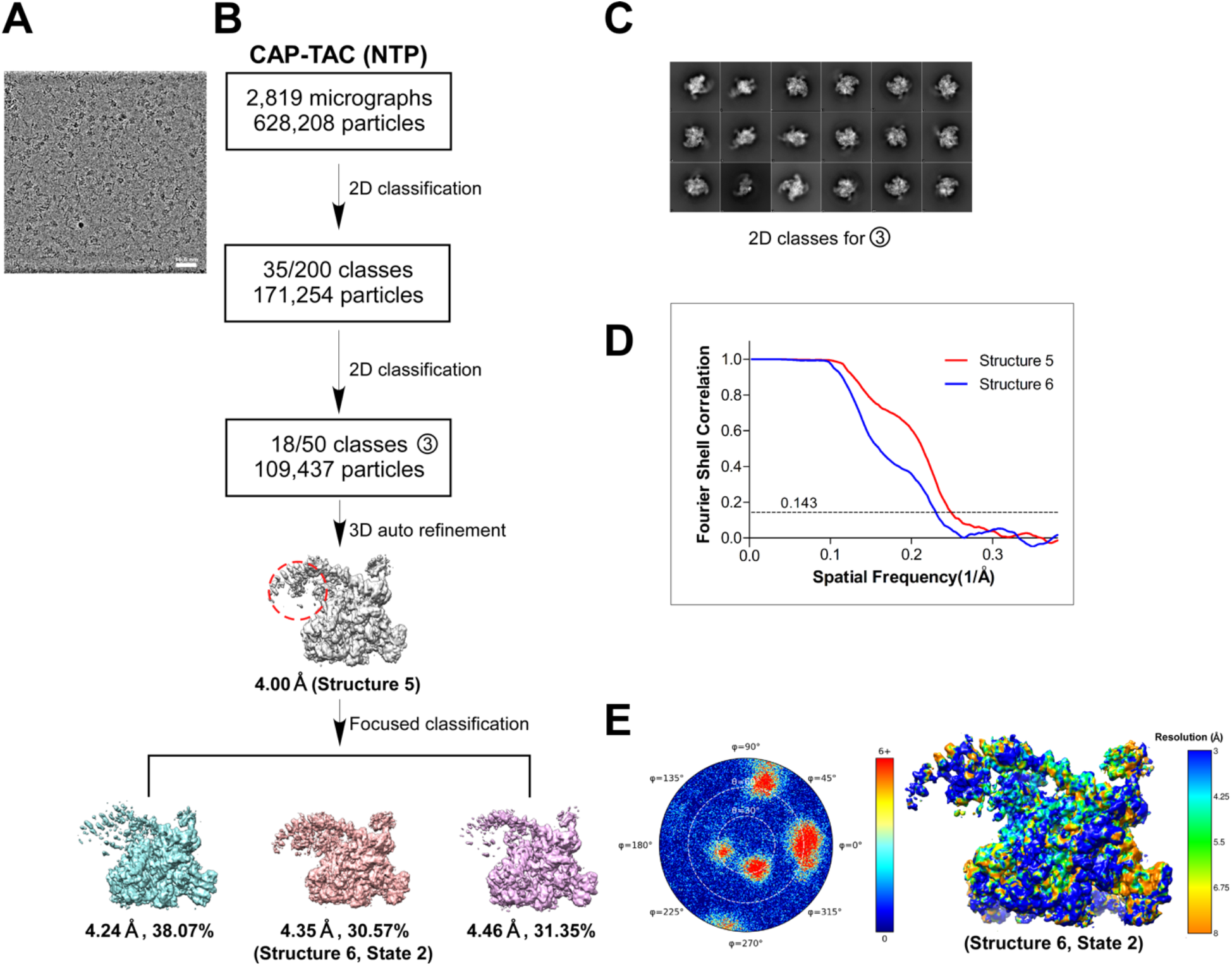
Cryo-EM images and data processing procedure for the CAP-TAC with *de novo* RNA transcript. A-C A representative micrograph (A), the flow chart of the cryo-EM image processing (see Methods) (B), and the selected 2D classes for structure 5 (C), respectively. D Gold-standard Fourier Shell Correlations (FSCs) of the maps for structures 5-6. E Angular orientation distribution (left) of the particles used in the final reconstruction and local resolution map (right) for structure 6 (state 2).

**Figure EV7.**
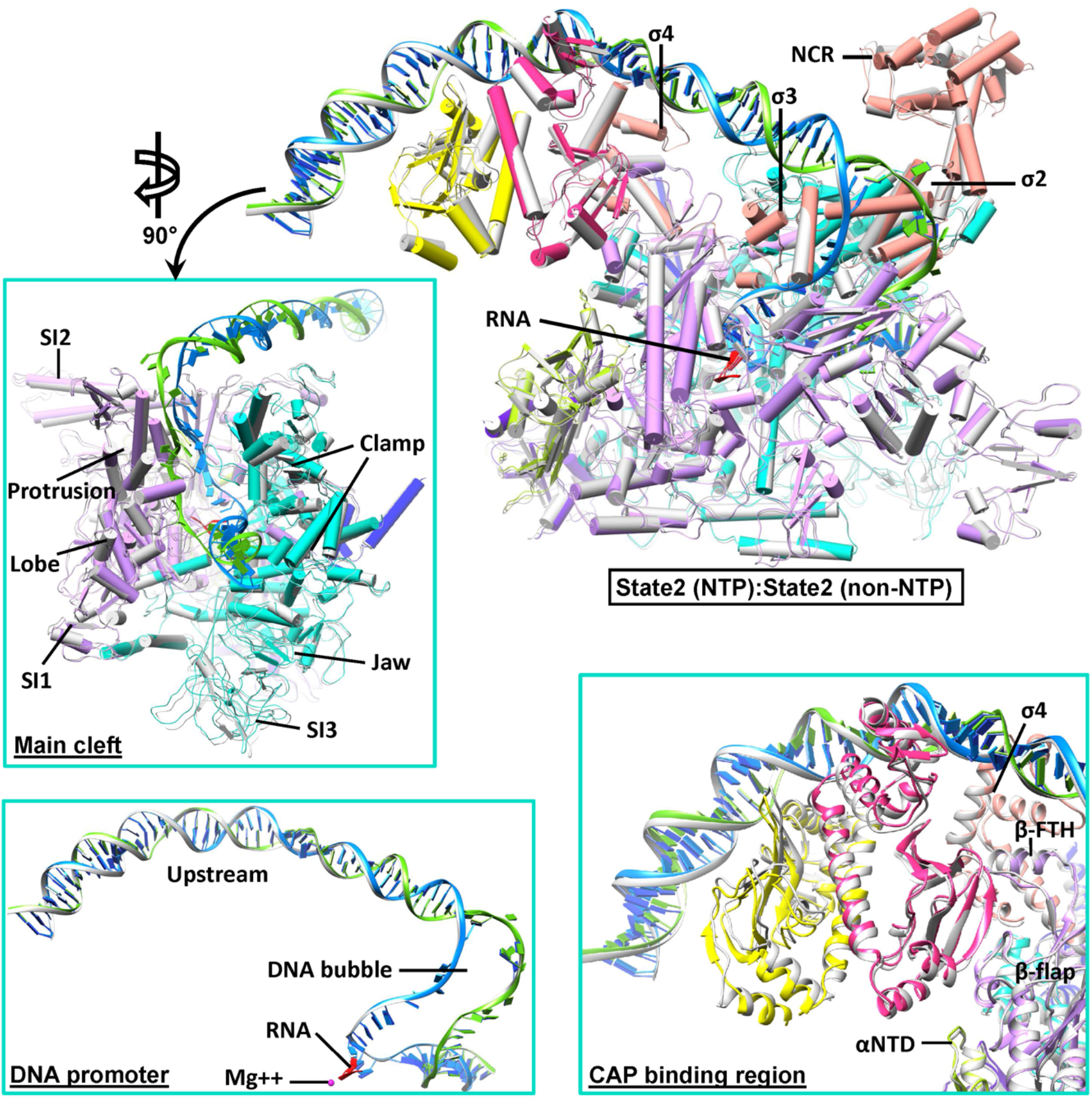
Superimposition between the two class-II CAP-TACs at the state 2. Superimposition of the state 2 CAP-TAC with *de novo* RNA transcript (colored) with the state 2 CAP-TAC without NTP incubation (gray) via the σ2 and σ3 domains of σ^70^ is shown. The color schemes are same as in Figure 1. The inserts are the close-up views of the superimpositions in the main cleft, the DNA promoter and the CAP binding region, which suggest the high structural similarity between them.

**Figure EV8.**
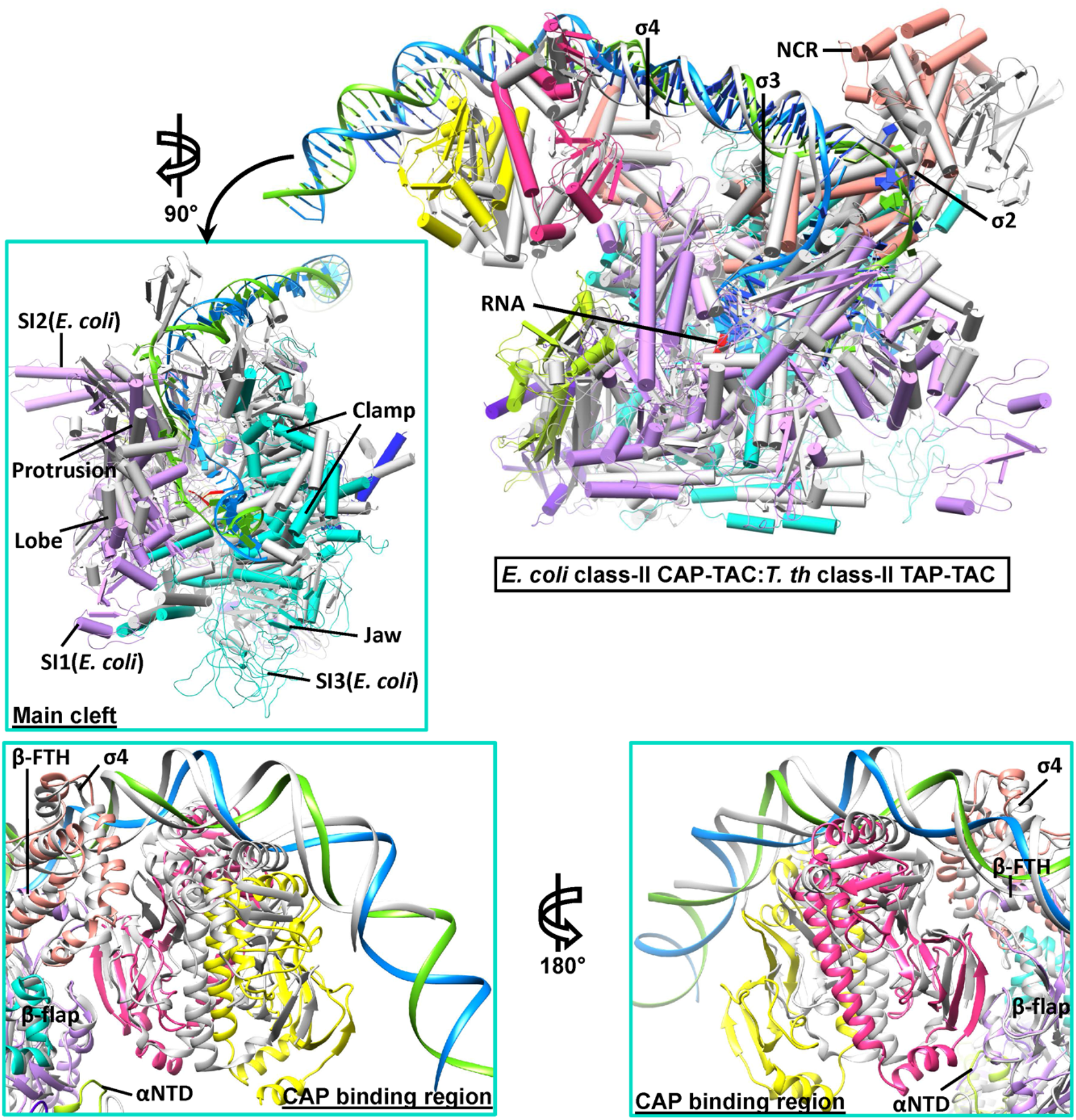
Superimposition between the *E. coli* and *T. th* class-II TACs. Superimposition of the *E. coli* class-II CAP-TAC (colored, state 2 with RNA) with the *T.th* class-II TAP-TAC (gray, PDB 5ID2) via the β subunit is shown. The color schemes are same as in Figure 1. The inserts are the close-up views of the superimpositions in the main cleft and the CAP binding region, suggesting the minor change in the width of main cleft, but apparent changes in the orientations of the domains on the interface between the CAP dimer and RNA holoenzyme.

**Table EV1.** Statistics of cryo-EM data collection, 3D reconstruction and model building.

| **Data collection/processing** | **CAP-TAC (non-NTP)** |  | **CAP-TAC (NTP)** |
| --- | --- | --- | --- |
| Microscope | Krios |  | Krios |
| Voltage (kV) | 300 |  | 300 |
| Camera | Falcon III |  | Falcon III |
| Camera mode | Counted |  | Counted |
| Defocus range ($\mu\text{m}$) | -0.8 ~ -2.6 | | -0.8 ~ -2.6 |
| Exposure time (s) | 45 |  | 36 |
| Dose rate ($e^-/\text{pixel/s}$) | 0.8 | | 0.8 |
| Magnified pixel size ($\text{\AA}$) | 0.9 | | 0.9 |
| **Reconstruction (focused maps)** | **State 1** | **State 2** | **State 2** |
| Software | cisTEM | cisTEM | cisTEM |
| Symmetry | C1 | C1 | C1 |
| Particles refined | 30296 | 37456 | 33455 |
| Resolution (auto-masked, $\text{\AA}$) | 4.52 | 4.29 | 4.35 |
| Access code | EMD-20287 | EMD-20288 | EMD-20286 |
| **Model Statistics** | **State 1** | **State 2** | **State2** |
| Number of residues (modeled) | 4245 | 4252 | 4255 |
| Map CC (Chimera) | 0.7382 | 0.8255 | 0.8314 |
| MolProbity score | 1.89 | 1.90 | 1.84 |
| All-atom Clashscore | 5.31 | 5.93 | 5.03 |
| Cβ deviations | 0 | 0 | 0 |
| Rotamer outliers | 0.23% | 0.11% | 0.00% |
| Ramachandran |  |  |  |
| Outliers | 0.69% | 0.29% | 0.42% |
| Favored | 87.64% | 89.06% | 89.19% |
| RMS deviations |  |  |  |
| Bond length | 0.008 | 0.007 | 0.006 |
| Bond angles | 1.221 | 1.053 | 0.969 |
| Access code | 6PB5 | 6PB6 | 6PB4 |

